## Supplementary Materials for "Improving few-shot learning-based protein engineering with evolutionary sampling" (from another version of this preprint: version 1, 2023-05-24)

### 1 Supplementary Methods and Results

#### 1.1 Training Data

An initial library of 34217 85aa (85 amino acid long) putative modulator peptides from diverse biological origins were experimentally screened for their ability to activate a synthetic genetic locus using the dCasMini Gene Expression Modulator System (dCasMini-GEMS) [1] in which the modulator peptide is fused to dCasMini and directed towards the synthetic genetic locus with gRNAs. In an independent follow-up screen, a subset of these sequences were re-tested, resulting in 173 sequences that we classified as validated gene activators ("positive hits"), giving a hit rate of $0.51\%$.

#### 1.2 Featurization

Since a set of peptide sequences is essentially composed of strings of 20 unique amino acid characters, each peptide sequence needs to be numerically featurized to be used as input to train supervised classification models. We explored several featurizations.

**OneHot Encoding** The simplest form of peptide featurization, OneHot encoding, transforms each of the 20 possible amino acid characters into a unique 20-dimensional vector with all values equal to zero except the amino acid index position with a 1. For our training set, this results in a list of 34217 arrays of size 85x20.

**ESM-2 Embedding** Evolutionary Scale Model 2 (ESM-2) is a transformer-based protein language model trained on over 60M sequences from the UniRef database [2, 3]. Training on such a huge diversity of protein sequences allows large protein language models (LPLMs) like ESM-2 to implicitly capture a variety of evolutionary, biochemical, and biophysical features that can be leveraged to improve functional predictions via transfer learning. In this study we use the 650M parameter 33-layer ESM-2 model which converts each 85aa peptide sequence into an array of size 85x1280. We additionally tested the larger 3B and 15B ESM-2 models but ultimately decided to use the 650M

*ZJ and RY contributed equally to this work.

Preprint. Under review.

parameter model embeddings since the lower parameter embedding performed comparably well. Furthermore, MHMCS and EMCS computational cost is dominated by the embeddings, therefore the smaller model enabled faster exploration.

#### 1.3 Model Training

OneHot and ESM-2 protein sequence embeddings were then used as input to train our two classification models: a decision-tree based model (XGBoost) [4], where we flatten the features by taking the mean [5], and a 1-dimensional convolutional neural network (CNN) [6]. Due to the extreme class imbalance, during all training we upsampled our positive hits so they accounted for 30% of the observations. All models were trained using using AWS Sagemaker Accelerated Computing P3 instances with NVIDIA V100 Tensor Core GPUs with PyTorch (v1.13). Model training results (PR-AUC and F1 scores) can be found in Supplementary Table 1 and Supplementary Table 2; for both XGBoost and CNN models, we observed a significant increase in PR-AUC and F1 scores with transfer learning using ESM-2 embeddings compared to OneHot encoding.

**XGBoost** We employed hyperparameter tuning with a grid search (parameters: "learning_rate", "n_estimators", "gamma", "subsample", "max_depth", "min_child_weight") and determined the top model based on PR-AUC score from 5-fold stratified cross-validation. XGBoost models were trained with tree_method="gpu_hist" and objective="binary:logistic". The top ESM-2 model included 500 trees with a max depth of 4 and was trained with parameters learning_rate=0.1, gamma=0, subsample=0.75, min_child_weight=0. The top model was then re-trained on the entire training set with upsampled positive hits and used as the XGBoost fitness function for exploring the fitness landscape.

**CNN** Model architecture for the 1-dimensional CNN models included 3 convolutional layers with rectified linear unit (ReLU) activation functions, with batch-normalization, dropout, and max-pooling, followed by two densely connected layers. We employed hyperparameter tuning with a grid search (parameters: "n_epochs", "weight_decay", "learning_rate", "batch_size", "dropout", "conv_size") and determined the top model based on F1 score from 5-fold stratified cross-validation. CNN models were trained using the AdamW optimizer with "BCEWithLogitLoss" loss function. The top ESM-2 model was trained for 30 epochs with a batch size of 16 using the AdamW optimizer with parameters learning_rate=1e-06 and weight_decay=0.05; this resulted in a network architecture with three convolutional layers containing 100 convolutional filters (kernel_size=10, stride=1) with batch-normalization and max-pooling followed by two densely-connected layers (200 neurons in first hidden layer and 10 neurons in second hidden layer). The top model was then re-trained on the entire training set with upsampled positive hits and used as the CNN fitness function for exploring the fitness landscape.

| Transfer Learning: Model Training (PR-AUC) | | | |
|---|---|---|---|
| Embedding | Fitness Model | PR-AUC (5-fold CV) | PR-AUC (30% test set) |
| OneHot | XGB | 0.487 | 0.549 |
| OneHot | CNN | 0.331 | 0.303 |
| ESM-2 | XGB | 0.573 | 0.629 |
| ESM-2 | CNN | 0.533 | 0.576 |

Supplementary Table 1: PR-AUC scores based on 5-fold cross-validation and testing on 30% holdout test set for top XGBoost and CNN models with OneHot and ESM-2 embeddings respectively.

| Transfer Learning: Model Training (F1 score) | | | |
|---|---|---|---|
| Embedding | Fitness Model | F1 score (5-fold CV) | F1 score (30% test set) |
| OneHot | XGB | 0.450 | 0.472 |
| OneHot | CNN | 0.457 | 0.432 |
| ESM-2 | XGB | 0.590 | 0.615 |
| ESM-2 | CNN | 0.562 | 0.519 |

Supplementary Table 2: F1 scores based on 5-fold cross-validation and testing on 30% holdout test set for top XGBoost and CNN models with OneHot and ESM-2 embeddings respectively.

#### 1.4 Sampling

All sampling runs were implemented on AWS Elastic Cloud Compute 2 g5.2xlarge instances with NVIDIA A10 Tensor Core GPU's. When using the ESM-2_650M parameter model to generate sequence embeddings, we found that the g5.2xlarge instances could evaluate roughly 50 independent 85aa sequences per second. We verified that the vast majority of the computational time was spent on sequence featurization.

**Metropolis Hastings Monte Carlo Search (MHMCS)** Algorithm 1 summarizes our implementation of MHMCS, with our choice of default parameters given in Supplementary Table 3.

**Algorithm 1** Metropolis Hastings Monte Carlo Search (MHMCS)

Initialize a peptide sequence $i$ and assign a temperature $T_i$.
Set minimum and maximum number of iterations $k_{min}$ and $k_{max}$, respectively.
Set maximum number of mutations $\mu$.
Set convergence condition $f(i) \geq C$ where $f(i)$ is the fitness of sequence $i$.
**repeat**
  Make random point mutations at $q$ loci for sequence $i$ to yield a new sequence that we call $j$, where $q \in \{1, \ldots, \mu\}$ is chosen uniformly at random
  Accept or reject sequence $j$ using the metropolis hasting criterion i.e. with probability $min(1, r_{mh})$, where $r_{mh} = \exp(\frac{f(j)-f(i)}{T})$
**until** $iterations > k_{max}$ **or** $((f_i \geq C)$ **and** $iterations > k_{min})$

**Default Sampling Parameters** Supplementary Table 3 summarizes the default parameters used for MHMCS and EMCS. Usually, the number of minimum and maximum iterations are decided by trying a few numbers and manually adjusting them to find a set of numbers where the algorithm is able to converge to a state with an acceptable fitness. This is done by simply plotting the fitness vs. the number of iterations. $T$ (also referred to as the temperature) decides how liberally non-optimal moves are accepted. A very high temperature will result in a very large proportion of non-optimal moves being accepted (>50-60%). This would make it almost impossible for the algorithm to converge, as the algorithm is unable to reject moves when it is close to an optimum. A smaller temperature makes the MHMCS algorithm more conservative, thereby only strictly accepting moves that increase the fitness or decrease it by a very small amount (favoring a higher resolution search). This makes it easier to move towards an optimum, but restricts the search space to directions where the fitness only increases. The selection of the temperature is therefore an important task, defining the tradeoff between the resolution (depth) and the width (area) of our search. EMCS allows us to search using different temperatures simultaneously. The temperatures were chosen by a combination of analyzing the fitness distribution of the training set (where the fitness scores spanned over multiple orders of magnitude between $10^{-4}$ and 1), and trial and error. The number of chains was set to 4 in order to achieve a good balance between fitness score coverage and computational cost. The algorithm was set to converge when one or more sequences were discovered with a fitness score $\geq 0.95$ and the maximum number of iterations was set to $10^5$.

| Default Sampling Parameters (MHMCS/EMCS) | | | | | | |
|---|---|---|---|---|---|---|
| Model | Number of Chains | Temperature | $\mu$ | $\gamma$ | $\alpha$ | $\beta$ |
| MHMCS | 1 | $2.5 \times 10^{-3}$ | 5 | - | - | - |
| MHMCS | 1 | $2.5 \times 10^{-4}$ | 5 | - | - | - |
| EMCS | 4 | $2.5 \times [10^{-4}, 10^{-3}, 10^{-2}, 10^{-1}]$ | 5 | 0.5 | 1 | 1 |

Supplementary Table 3: Default sampling parameters for EMCS and MHMCS. Key (as defined in main text): $\mu$ (Max. Mutation Events Per Iteration), $\gamma$ (Crossover Rate), $\alpha$ (Max. Crossover Events Per Iteration), $\beta$ (Max. Swap Events Per Iteration)

**Ensemble Model** When sampling using the ensemble model, each new proposed sequence was evaluated using both XGBoost and CNN models, where fitness was defined to be the average of the respective fitness scores of the XGBoost and CNN models. This 'combined' fitness was then used to accept or reject any proposed moves.

**Crossover Criterion** The crossover criterion ($min(1, r_c)$) is used to determine whether the set of new sequences $i_2, j_2$ generated by crossing over two sequences $i_1, j_1$ are accepted or rejected. Here, $r_c$ is defined as:

$$r_c = \exp\left(\frac{f(i_2) - f(i_1)}{T_i} - \frac{f(j_2) - f(j_1)}{T_j}\right) \tag{1}$$

With the condition that $T_i \leq T_j$, and the sequences $i_2$, $j_2$, and $i_1$, $j_1$ are ordered such that $f(i_1) \geq f(j_1)$ and $f(i_2) \geq f(j_2)$. The proposed sequences $i_2$, $j_2$ are accepted with probability $min(1, r_c)$ and assigned to temperatures $T_i$ and $T_j$ respectively.

By rearranging the terms in $r_c$, we can see that $r_c$ will always be greater than 1 if

$$\frac{i_2}{T_i} - \frac{j_2}{T_j} > \frac{i_1}{T_i} - \frac{j_1}{T_j} \tag{2}$$

This condition always holds true for optimal moves. Including the move in which the fitness of both sequences improves as in $f(i_1) < f(i_2)$ and $f(j_1) < f(j_2)$, thereby moving both sequences to a region with a higher fitness landscape. If $f(i_1) < f(i_2)$ but $f(j_1) > f(j_2)$, that is, the fitness of one sequence improves while the fitness of of the other sequence is worsened, then the proposed move is only guaranteed to be accepted if the condition in Supplementary Equation 2 holds. If the condition in Supplementary Equation 2 does not hold, then the proposed move is sub-optimal and is only accepted with probability equal to that defined in Supplementary Equation 1. In general, the likelihood of acceptance of a sub-optimal moves rapidly decreases as the temperature difference $T_j - T_i$ increases, thereby suggesting that a large temperature difference penalizes sub-optimal moves more than a smaller temperature difference.

#### 1.5 Entropy Calculations

The change in entropy for a given iteration gives us a measure of the change in information observed for that iteration. Figure 3a shows the entropy change distribution of two simulated MHMCS and EMCS runs under default parameters. However, the number of iterations was preset to $10^7$ to average over a sufficiently large number of iterations (for reference, Supplementary Table 8 shows the convergence times of a general run under default parameters to be $< 10^4$).

The information entropy of a given sequence is defined as:

$$S = -\sum_i p_i log(p_i), \tag{3}$$

where $i$ runs over the 20 possible amino acid residues, and $p_i$ is the ratio of the count of a given amino acid residue divided by the length of the peptide. For example, a peptide sequence of 'AAASST' would have $p_i$ equal to $\frac{3}{6}$, $\frac{2}{6}$, $\frac{1}{6}$ for residues 'A', 'S', and 'T' respectively, and zero for all other residues. For MHMCS, the entropy change for a given iteration is defined as $\Delta S = S_{proposed} - S_{initial}$. Where $S_{initial}$ is the entropy of the sequence at the beginning of the iteration, i.e. prior to the application of a mutation operator.

For EMCS, the entropy change is first independently calculated for each of the chains ($N_{chains} = 4$ for default parameters). Then, the value of the maximum entropy change is retained and recorded for a given iteration. It is important to note that the entropy changes in each of the chains occur due to the application of the mutation and/or the crossover operator, which in turn depends on the choice of parameters we use. In Figure 3a, we only show this difference for the default parameters given in Supplementary Table 3. Supplementary Table 4 provides additional data for Fig. 3a.

| Entropy Change Distribution (MHMCS/EMCS) for $10^7$ iterations using default parameters (Fig.3a) | | | | | |
|---|---|---|---|---|---|
| Model | Mean | Standard Deviation | $25^{th}$ | $50^{th}$ | $75^{th}$ |
| EMCS | 0.334 | 0.229 | 0.146 | 0.252 | 0.437 |
| MHMCS | 0.092 | 0.101 | 0.013 | 0.061 | 0.133 |

Supplementary Table 4: Additional data for Figure 3a

#### 1.6 Sequence Diversity

To ensure that our sequences were sufficiently different from the training data when initialized from known positive sequences, we calculated the minimum edit (Levenshtein) distance of all high fitness sequences to the full set of sequences in the training dataset, thereby giving us a measure of sequence diversity. Supplementary Figure 1 summarizes this result. We note that, in general, EMCS yields sufficiently diverse sequences compared to MHMCS when starting from positive examples. While high temperature MHMCS runs yielded slightly more diverse sequences, it also failed to converge the majority of the time (275/500 runs).

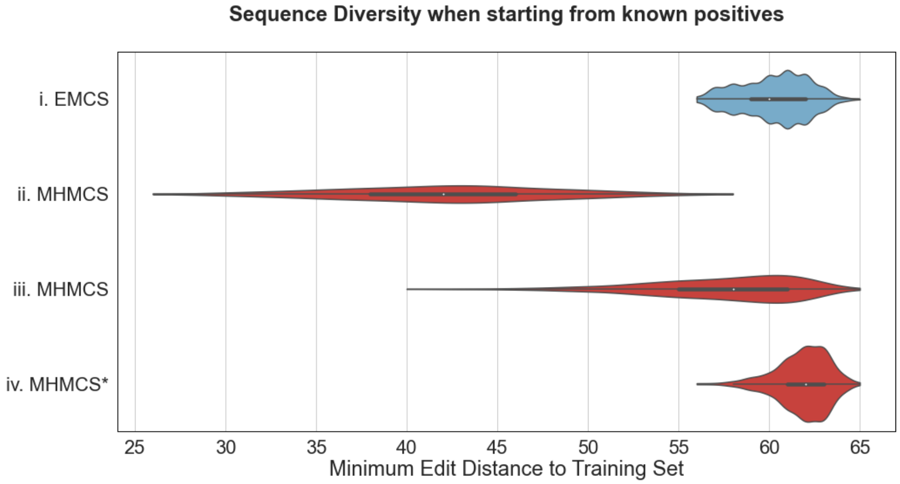


Supplementary Figure 1: Minimum edit distance of converged sequences to the original training dataset initialized from known positive hits when using i) EMCS under default parameters (N = 2000), ii) MHMCS with $T = 2.5 \times 10^{-4}$ (N = 500), iii) MHMCS with $T = 2.5 \times 10^{-3}$ (N = 500), and iv) $T = 2.5 \times 10^{-2}$ (N = 500). Although the average edit distance of the sequences obtained with iv) was slightly higher than i), the failure rate at iv) exceeded 50%. Refer to Table S9 for additional data.

#### 1.7 Experimental Screening

**Library Synthesis** For experimental validation, we designed a library of 4600 novel peptide sequences (and included 300 previously validated negative control peptides with random sequences) based on our exploration of the modulator fitness landscape (Supplementary Table 5). The 85aa peptide sequences were reverse-translated into 255bp nucleotide sequences using randomized codons to minimize GC bias [7], which were then synthesized as a pooled library by Twist Bioscience.

**Wetlab Screening** The library was cloned downstream of dCasMINI in a lentiviral expression plasmid at high coverage ($\approx$ 300x) and packaged into lentiviral particles (2 lentivirus batches were produced by transfecting separate dishes of 293T cells). K562 cells bearing a GFP reporter were infected in quadruplicate (2 biological replicates receiving different lentiviral batches, each with 2 technical replicates) at 1000x coverage with low multiplicity of infection (MOI = 0.3) to ensure that each infected cell received only one dCasMINI-modulator fusion construct. Infected cells were then enriched by Puromycin selection. At 7 days post-transduction, cells were sorted using fluorescence-activated cell sorting (FACS) to isolate GFP_OFF and GFP_ON populations (Supplementary Figure 2). For each replicate, $\approx 10^6$ GFP_ON cells were harvested. Genomic

| Validation library composition | | |
|---|---|---|
| Algorithm | Initialization | Total Sequences |
| EMCS | Known | 1310 |
| EMCS | Random | 1290 |
| MHMCS | Random | 2000 |
| Negative Controls | n/a | 300 |

Supplementary Table 5: Overall composition of peptides in validation library sorted by choice of sampling algorithm. Initialization column denotes the sampling algorithm's starting sequence as either randomly initialized ("Random"), or known positive hit ("Known").

DNA was prepared from GFP_ON, GFP_OFF, and unsorted populations, and barcoded modulator sequences were amplified using PCR primers to add Illumina i5 and i7 indexed adapter sequences. Pooled libraries were then sequenced on an Illumina NextSeq (Gladstone Genomics Core).

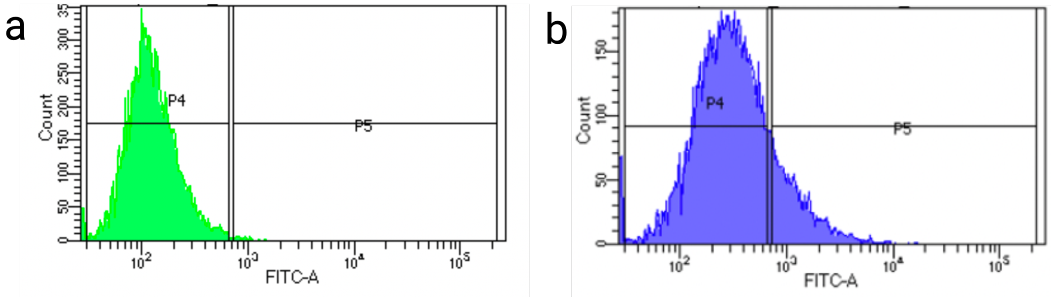


Supplementary Figure 2: Representative FACS histograms illustrating cell counts within GFP_OFF (P4) and GFP_ON (P5) gates in un-infected cells (a) and cells infected with the validation library (b). FITC-A is a readout for GFP levels.

**Bioinformatic Analysis** Raw sequencing reads were first aligned to a list of DNA barcodes (each putative modulator peptide is associated with a unique 12bp DNA barcode) in order to generate a count matrix. Read counts of technical replicates were then summed by barcode resulting in 2 biological replicates of GFP_ON and GFP_OFF libraries respectively. DESEQ2 [8] was used to determine which peptides were differentially enriched in the GFP_ON bins vs the GFP_OFF bins resulting in 357 positive hits (FDR<0.05 and log2FC>0) that were capable of activating the synthetic gene locus ((Supplementary Figure 3a, Supplementary Table 6, Supplementary Table 7). As expected, negative controls were depleted from the GFP_ON bin as these randomly encoded peptides were unable to activate the synthetic fluorescent reporter (Supplementary Figure 3b).

| Experimental Results: Sampling Algorithm | | | | |
|---|---|---|---|---|
| Algorithm | Initialization | Total Sequences | Number of Hits | Hit Percentage |
| EMCS | All | 2600 | 338 | 13% |
| MHMCS | All | 2000 | 18 | 0.9% |
| EMCS | Known | 1310 | 270 | 20.6% |
| EMCS | Random | 1290 | 68 | 5.3% |
| MHMCS | Random | 2000 | 18 | 0.9% |
| Negative Controls | n/a | 300 | 1 | 0.33% |

Supplementary Table 6: Final hit percentage of novel sequences sorted by choice of sampling algorithm. Initialization: Notes the sampling algorithm starting sequence as either randomly initialized, or known positive. Total Sequences: Number of sequences that were identified as high fitness by the machine learning model. Number of Hits: Number of sequences that validated experimentally.

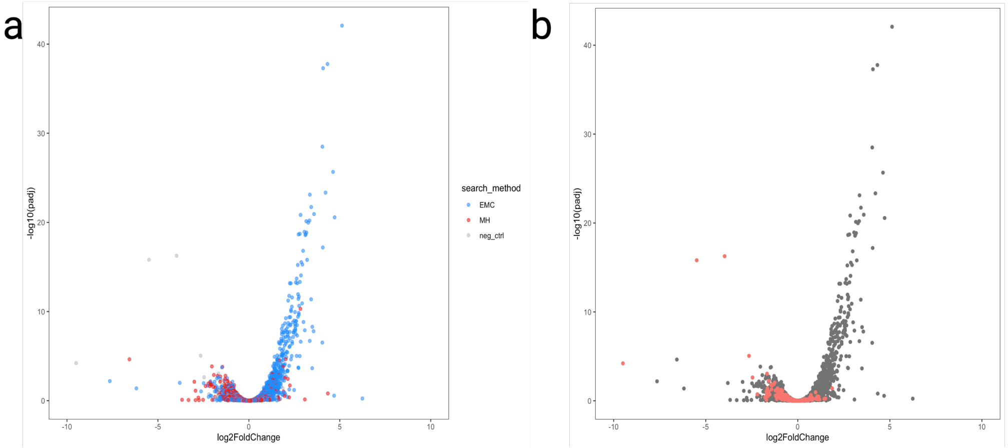


Supplementary Figure 3: Volcano plots illustrating statistical signficance (adjusted p-value) and activation strength (Log2FoldChange) of experimentally screened modulator peptides colored by sampling method (a) and highlighting negative controls (b).

| Experimental Results: Model Choice | | | |
|---|---|---|---|
| Fitness Model | Total Sequences | Number of Hits | Hit Percentage |
| Ensemble (XGB+CNN) | 1000 | 135 | 13.5% |
| XGB | 1800 | 145 | 8.06% |
| CNN | 1800 | 76 | 4.2% |
| Negative Controls | 300 | 1 | 0.33% |

Supplementary Table 7: Final hit percentage of novel sequences sorted by choice of machine learning model. Total Sequences: Number of sequences that were identified as high fitness by the machine learning model. Number of Hits: Number of sequences that validated experimentally.

#### 1.8 Principal Component Analysis (PCA)

In order to visualize how the library of 4600 novel proposed sequences compared to the original training data, we performed principal component analysis (PCA) with Scikit-learn in Python (v3.9). Briefly, we combined peptide sequences from our trianing data with the 4600 sequences in the validation library, represented each sequence with OneHot encoding and then flattened the encoding by taking the mean. Features were scaled using StandardScaler() prior to calculating and visualizing principal components.

#### 1.9 Additional Data

| Additional Data: Number of Iterations to Convergence (Figure 3b) | | | | | | | | |
|---|---|---|---|---|---|---|---|---|
| Model | Inclusion Criteria | Number of Runs | Number of Sequences | Mean | Standard Deviation | $25^{th}$ | $50^{th}$ | $75^{th}$ |
| i. MHMCS | $f > 0.95$ | 2571 | 2361 | 8837 | 14409 | 1509 | 3501 | 8892 |
| ii. EMCS | $f > 0.95$ | 1171 | 1171 | 3950 | 4172 | 1500 | 2825 | 4985 |
| iii. MHMCS | $f > 0.5$ | 2571 | 2571 | 16284 | 28545 | 1630 | 3981 | 12693 |
| iv. EMCS | $f > 0.5$ | 1171 | 2720 | 2453 | 3822 | 35 | 1075 | 3546 |

Supplementary Table 8: Additional data for Figure 3b

| Additional Data: Sequence Diversity measured by edit distance when starting from known positives | | | | | | |
|---|---|---|---|---|---|---|
| Model | Number of Sequences | Mean | Standard Deviation | $25^{th}$ | $50^{th}$ | $75^{th}$ |
| i. EMCS | 2000 | 60.242 | 1.89 | 59 | 60 | 62 |
| ii. MHMCS | 500 | 49.21 | 5.96 | 38 | 42 | 46 |
| iii. MHMCS | 500 | 57.4 | 4.21 | 55 | 58 | 61 |
| iv. MHMCS* | 225 | 61.82 | 1.43 | 61 | 62 | 63 |

Supplementary Table 9: Additional data for Figure S1

### References


[1] Giovanni Carosso, Robin W Yeo, T Blair Gainous, M Zaki Jawaid, J Luke Campos, Xiao Yang, Vincent Cutillas, Stanley L Qi, Timothy P Daley, and Daniel Hart. Discovery and engineering of hypercompact transcriptional modulators for robust and durable target gene activation. *In Preparation*, 2023.

[2] Zeming Lin, Halil Akin, Roshan Rao, Brian Hie, Zhongkai Zhu, Wenting Lu, Nikita Smetanin, Robert Verkuil, Ori Kabeli, Yaniv Shmueli, Allan dos Santos Costa, Maryam Fazel-Zarandi, Tom Sercu, Salvatore Candido, and Alexander Rives. Evolutionary-scale prediction of atomic-level protein structure with a language model. *Science*, 379(6637):1123–1130, March 2023. doi: 10.1126/science.ade2574. URL `https://doi.org/10.1126/science.ade2574`.

[3] Alexander Rives, Joshua Meier, Tom Sercu, Siddharth Goyal, Zeming Lin, Jason Liu, Demi Guo, Myle Ott, C. Lawrence Zitnick, Jerry Ma, and Rob Fergus. Biological structure and function emerge from scaling unsupervised learning to 250 million protein sequences. *Proceedings of the National Academy of Sciences*, 118(15), April 2021. doi: 10.1073/pnas.2016239118. URL `https://doi.org/10.1073/pnas.2016239118`.

[4] Tianqi Chen and Carlos Guestrin. XGBoost: A scalable tree boosting system. In *Proceedings of the 22nd ACM SIGKDD International Conference on Knowledge Discovery and Data Mining*, KDD '16, pages 785–794, New York, NY, USA, 2016. ACM. ISBN 978-1-4503-4232-2. doi: 10.1145/2939672.2939785. URL `http://doi.acm.org/10.1145/2939672.2939785`.

[5] Sanjeev Arora, Yingyu Liang, and Tengyu Ma. A simple but tough-to-beat baseline for sentence embeddings. In *International conference on learning representations*, 2017.

[6] Yann LeCun, Yoshua Bengio, and Geoffrey Hinton. Deep learning. *Nature*, 521(7553):436–444, May 2015. doi: 10.1038/nature14539. URL `https://doi.org/10.1038/nature14539`.

[7] Valentin Zulkower and Susan Rosser. DNA chisel, a versatile sequence optimizer. *Bioinformatics*, 36 (16):4508–4509, July 2020. doi: 10.1093/bioinformatics/btaa558. URL `https://doi.org/10.1093/bioinformatics/btaa558`.

[8] Michael I Love, Wolfgang Huber, and Simon Anders. Moderated estimation of fold change and dispersion for RNA-seq data with DESeq2. *Genome Biology*, 15(12), December 2014. doi: 10.1186/s13059-014-0550-8. URL `https://doi.org/10.1186/s13059-014-0550-8`.
